# Environmental sources of bacteria and genetic variation in behavior influence host-associated microbiota

**DOI:** 10.1101/296228

**Authors:** Alexandra A. Mushegian, Roberto Arbore, Jean-Claude Walser, Dieter Ebert

## Abstract

In many organisms, host-associated microbial communities are acquired horizontally after birth. This process is believed to be shaped by a combination of environmental and host genetic factors. We examined whether genetic variation in animal behavior could affect the composition of the animal’s microbiota in different environments. The freshwater crustacean *Daphnia magna* is primarily planktonic, but exhibits variation in the degree to which it browses in benthic sediments. We performed an experiment with clonal lines of *D. magna* showing different levels of sediment-browsing intensity exposed to either bacteria-rich or bacteria-poor sediment or whose access to sediments was prevented. We find that the bacterial composition of the environment and genotype-specific browsing intensity together influence the diversity and composition of the *Daphnia*-associated bacterial community. Exposure to more diverse bacteria did not lead to a more diverse microbiome, but greater abundances of environment-specific bacteria were found associated with host genotypes that exhibited greater browsing behavior. Our results indicate that individual behavior can mediate genotype-by-environment interaction effects on microbiome composition.

**Summary statement:** Genetic differences in Daphnia behavior contribute to the amount of environmental bacteria present in their microbiome

## Introduction

Every multicellular organism is colonized by a community of microorganisms: its microbiota (1). The host provides a habitat for a complex and dynamic consortium of microorganisms, many of which have fundamental influences on the host’s well-being. A central concern in both infectious disease epidemiology and in studies of host-associated microbial community ecology is the transmission of microbes between host individuals and between hosts and the environment. Many bacterial assemblages are transmitted from host mother to offspring (2) or within social groups (3), but the diversity of microbiota typically changes over time depending on the microbes available in the environment (4, 5). In some cases, environmentally acquired microbes are even essential for the completion of postembryonic development (6, 7). Thus microbes from the environment can be co-opted as part of the microbiota, or can affect host health during a transient occupation (8).

Environmental effects on microbiota community structure have been extensively documented (9, 10) and studies on model organisms have started to shed light on the relative importance of environmental and host genetic factors in determining microbiota composition (11, 12). Recently, the focus has been moving towards a better understanding of the mechanisms of bacterial aqusition from the environment. Host genetics have been shown to play a role in the establishment of microbial associations through microbial recognition, immune selection, and determination of the biochemical niche (12). Importantly, these processes select microbes after the host has come in contact with bacterial communities in the environment. The initial encounter may be a key phase of the host’s colonization by microbes. If host genetics influence interaction with the environment, for example through the expression of behavioral variation, it may influence the initial encounters with environmental bacteria and thus affect the composition of the host microbiota.

Many animals utilize different habitats according to behavioral strategies collectively termed habitat selection. If habitats differ in their microbial communities, host behavior influencing habitat choice and the microbiome may influence each other. Hosts may have evolved strategies to ensure or avoid encounter with beneficial and pathogenic microorganisms. Avoidance behaviors of harmful bacteria are well documented, and behavior is considered one of the first lines of defense against infectious disease. For example, the nematode *Caenorhabditis elegans* actively avoids pathogenic bacteria and the genetic determinants of this behavior have been worked out (13). The opposite case, where a host’s behavior is involved in the acquisition of beneficial bacteria from the environment, has received less attention, despite speculation about the role of human behaviors such as outdoor play in preventing autoimmune diseases (14). The overall effects of host habitat choice behavior on microbiome composition have not, to our knowledge, been explored in any system. An analysis of natural genetic variation in behavioral traits potentially influencing microbiota acquisition is therefore timely (15). If variation in behavior affects the composition of the host’s microbial community, then behavior could underlie some genotype-environment interaction effects on microbiota. The goal of this study was to examine the effect of genetic variation in host behavior on microbiota composition in different environments using the freshwater planktonic crustacean *Daphnia magna*.

Recently, it has been shown that *D. magna* microbiota play a major role in host fitness (16), that both host clonal line and environmental factors are determinants of microbiota community structure (17) and that genotype-specific microbiomes can mediate daphnids’ adaptive traits (18). However, little is known about the mechanisms by which the host acquires microbiota from the environment. A specific behavior, termed sediment browsing, mediates the interaction between *D. magna* and bottom sediments of ponds and lakes (19, 20). During browsing, the animals swim along the sediment surface, stirring up particles, and then ingest the particles by filter feeding. Besides representing valuable food reservoirs, sediments are likely important environmental sources of bacteria. Therefore, the physical contact with the sediments resulting from browsing might present both disease risks and benefits from increased contact with bacteria. Previous work found evidence of genetic variation for the levels of browsing activity in *D. magna* (21).

We performed a laboratory experiment in which we analyzed the browsing behavior and microbiota of 12 genetically distinct *D. magna* clones allowed to browse in sediment. The animals were exposed to three different treatment conditions, where they had access to either previously autoclaved (i) or untreated (“natural” and therefore microbe-rich) sediments (ii), or where their access to natural sediment was prevented (iii) (Figure 1A). We hypothesized that *D. magna* clones exhibiting more intense browsing behavior would have more diverse microbiota in conditions where they had access to bacteria-rich sediment, whereas the microbiome would be less affected by the bacterial environment in genotypes that browsed less. In this experiment, we made no assumptions about whether bacteria found in the sediments were beneficial, harmful, or neutral for the host, nor whether they colonized *Daphnia* stably or transiently; therefore, the patterns observed here could be applicable to studies of disease, microbiota, or general environmental microbial community dynamics. Our analysis illustrates how a behavioral trait can mediate the interplay between genetic and environmental variation in the establishment of host-microbe associations.

**Figure 1:**
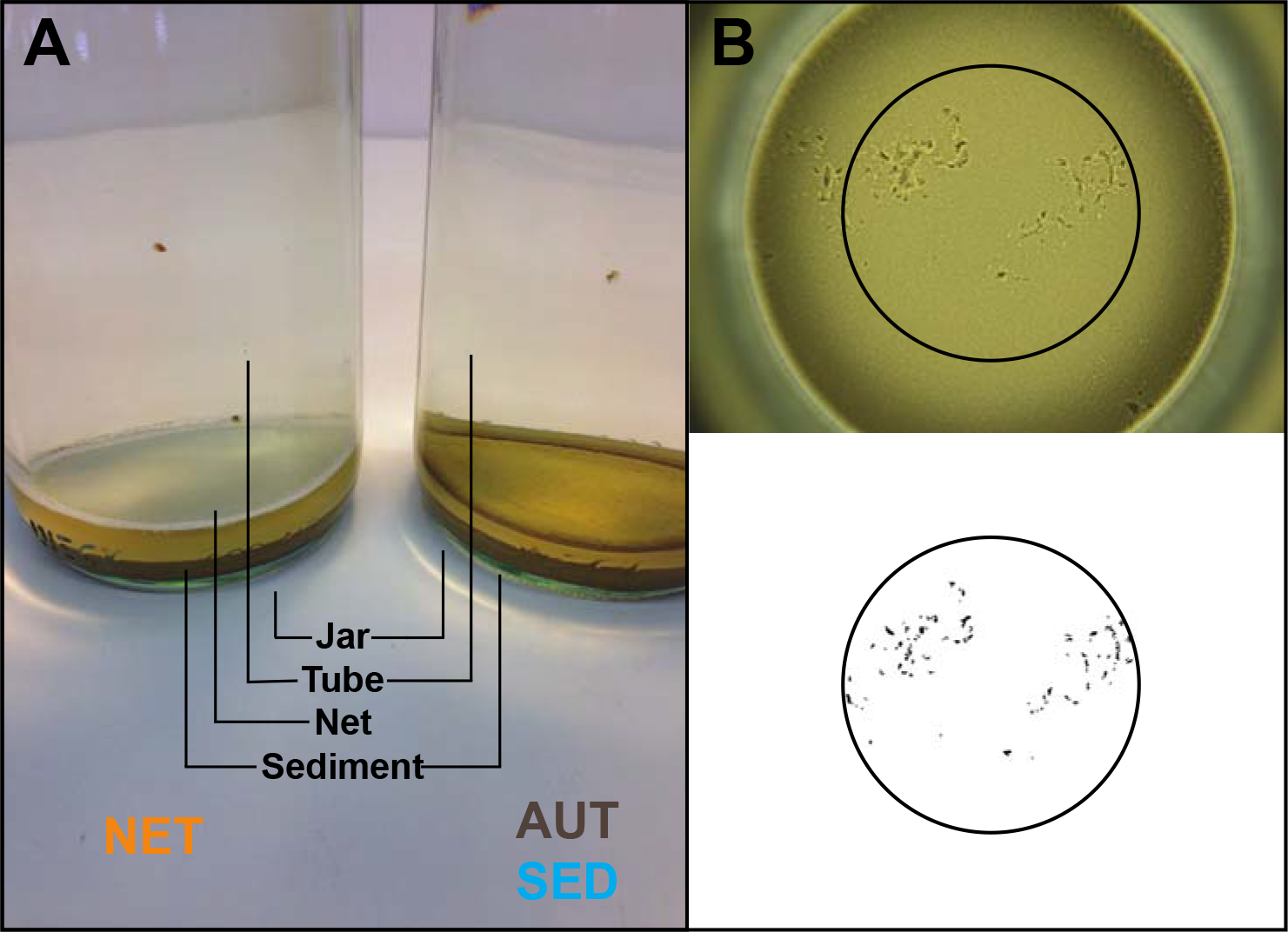
Experimental set-up and browsing behavior assay. A: the jars used in the experiment had a bottom layer of fine loess and contained two animals each; the animals were prevented from browsing on untreated sediments by a net placed 5 mm above the sediment surface (NET, right) or were allowed to browse on autoclaved sediments (AUT) or untreated sediments (SED) (left). B: traces left by one animal browsing on a sediment surface for 30 minutes and the same picture after processing for quantification of the browsing behavior.

## Methods

### Overview of the experiment

In this study, we combined the analysis of animals with constitutive (genetic) differences in browsing behavior with manipulations of the environment that affected animals’ access to the sediments. Animals were either exposed to natural sediment, to autoclaved sediment, or to natural sediment blocked by a permeable net barrier (Figure 1A). In order to analyze both the browsing behavior of the animals and their microbiota, we placed two animals in each jar; of these pairs of animals, after 6 days of exposure to the different treatments, one animal was used to assay browsing behavior while the other was used for microbiota analyses.

### Experimental animals

*D. magna* reproduces by cyclical parthenogenesis. Clonal populations can be generated and propagated in the laboratory through asexual reproduction. Here, we refer to such genetically identical individuals as “replicates” or “animals” while we refer to different genetic lines as “clones.” In this study we used 12 *D. magna* clones from our stock collection, originating from different populations (Table 1). The animals were propagated from stock cultures maintained in the laboratory in standardized conditions and without any effort to modify their microbiota. The browsing behavior of these clones has been assessed before (21) and was shown to differ among genotypes.

**Table 1.**
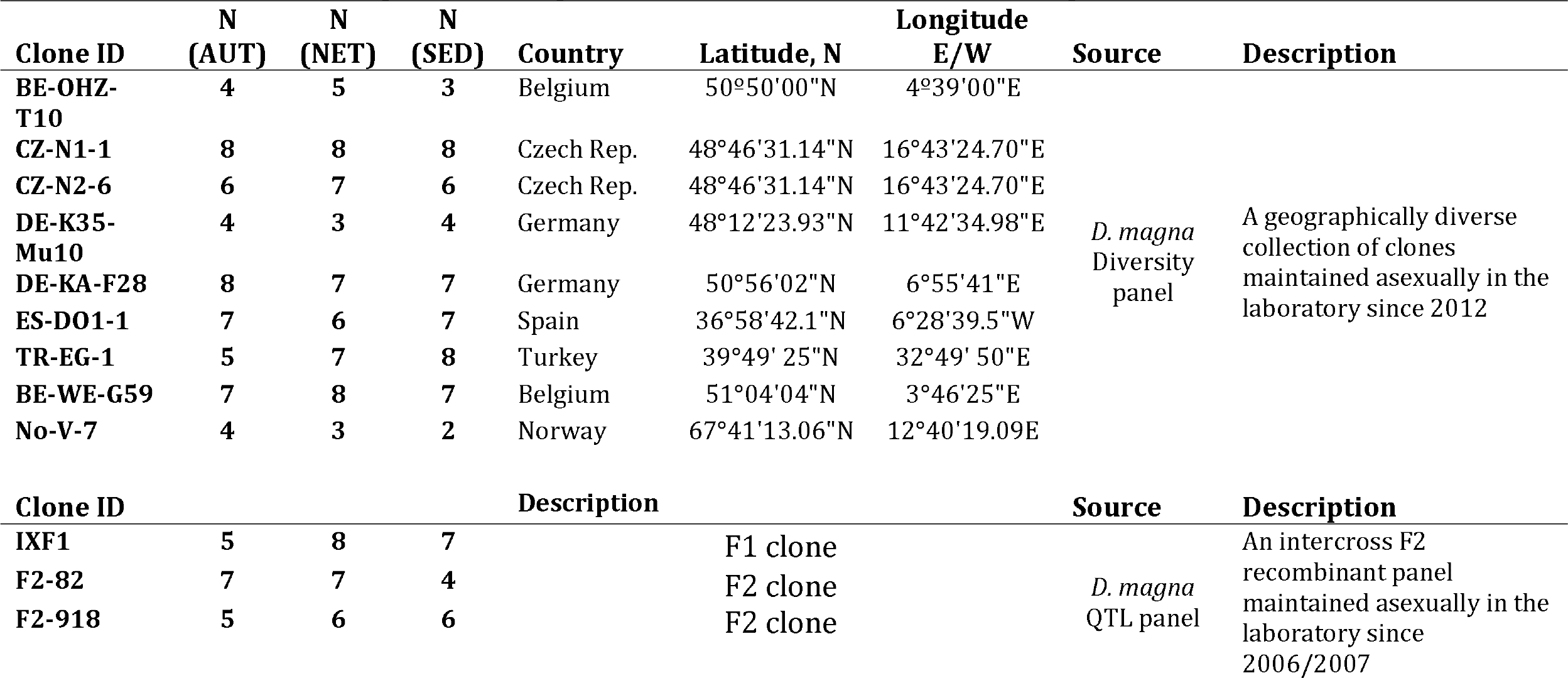
Names, number of individual replicates included in the microbiota analyses in the three treatments (AUT, NET and SED) and origin information of the 12 *Daphnia magna* clones used in this study. AUT: Exposure to autoclaved sediment; NET: prevented exposure to untreated sediment; SED: exposure to untreated sediments.

All animals used in this study were females. Prior to the experiment, every clone was propagated in individual replicates for three generations in order to minimize variation due to maternal effects. These animals were kept individually in 100-ml glass jars filled with 80 ml of ADaM (*Daphnia* medium (22)) randomly distributed within trays in incubators with a 16:8 light/dark cycle and constant temperature of 20 °C. To establish every generation, the animals were isolated at 4 days old and fed daily with chemostat-grown green algae *Scenedesmus* sp: 1 × 10^6^ algae cells/animal until day 5, 2 × 10^6^ until day 8, 2.5 × 10^6^ until day 10, 3 × 10^6^ until day 12, and 5 × 10^6^ onwards. The animals were transferred to fresh medium when they were 12 days old and thereafter every day.

For the experiment, we used animals from the 4^th^ generation of each of the 12 clones. These animals were kept in groups of 8 siblings belonging to one clutch of one mother. At 4 days old (± 1 day), 6 animals from every clutch were randomly assigned in pairs to individual jars divided into the three different treatments (split brood design); each such jar containing a pair of animals was an experimental replicate. In total, we included in the experiment 540 animals (270 pairs) corresponding to 15 pairs of clone BE-OHZ-TIO, 18 pairs of clones DE-K35-MulO and NO-V-7, 21 pairs of clone F2-918, and 24 pairs of each of the remaining clones. Variation in replicate numbers resulted from differences in availability of female offspring at the time that the treatments were established.

### Experimental design

The experiment was conducted in cylindrical glass jars (height = 20 cm; diameter = 6.5 cm) (Fig. 1A) kept in cardboard boxes on shelves in a climate room (16:8 light/dark cycle at 20 °C), loosely covered with transparent plastic film and top-illuminated with neon lights. In this way, light only entered the jars from the top. All the experimental jars were first filled with 400 ml of medium. 15 ml of a suspension of loess (fine silt) was then carefully deposited on the bottom using a serological pipette. The loess was previously collected from a soil stock, suspended in water, passed through a 200 μm filter and washed to remove very fine particles. After two days of sedimentation, the loess formed a 1 cm layer at the bottom of the jar. Then, an acrylic tube (height = 21 cm; diameter = 5 cm) was inserted into the jars and kept in position with a plastic ring fixed to the opening of the jar, so that its lower end was positioned close to the sediment surface. In one treatment (NET), the acrylic tube was closed with a 500 μm net at the lower end (suspended 5 mm above the sediment surface) preventing animals from direct contact with the sediment (Fig.1A left). In the other two treatments (AUT and SED), the acrylic tubes had no net so that the animals had free access to the sediment (Fig.1A right). In the AUT treatment, the loess was previously autoclaved while in the SED and the NET treatment the loess was left untreated (“natural”). (After autoclaving, AUT sediment was handled in the same way as natural sediment, i.e. exposed to nonsterile media and laboratory environment.) After inserting the tubes, the jars were left undisturbed for two days before the animals were introduced in order to allow the sediment to settle. Immediately before the experiment, the sediments of three jars of each of the SED and the AUT treatment were sampled and frozen at −20 °C; these sampled jars were not used further.

Two animals from the same clutch were carefully introduced into the inner tube of each jar. The 264 jars, each containing one pair of animals, were evenly distributed among the treatments and their positions in the incubator room were randomized. The animals remained in the experimental jars for 6 days. During this time, the animals were carefully fed twice daily with 2.5 × 10^6^ algal cells. At day 6, all animals were collected and one member of every pair was assigned to the behavioral assay (see below) and the other was frozen for later DNA extraction. 32 pairs of animals were lost or damaged during the experiment and were excluded from further analyses. At the end of the experiment, 3 sediment samples from the NET treatment and 3 sediment samples from the AUT treatment were collected and frozen at −20°C.

### Behavioral analysis

The animals for the behavioral assay were transferred individually from the sediment jars to 100-ml glass jars filled with medium and kept in an incubator and fed daily with 5 × 10^6^ algal cells. The behavior assay was conducted over two days when the animals were 12 to 14 days old with all replicates for the different clone by treatment combinations evenly distributed across time. The behavior assay was performed as described previously (21). Briefly, we measured the traces left by individual replicate animals on a smooth surface layer of sediment (loess) at the bottom of tall cylindrical glass jars (20 cm tall, 6.5 cm diameter; Fig. 1B) during 30 minutes. The sediment surface was photographed before animals were released (time 0), using a ring light to ensure uniform illumination. The jar was then transferred into a cardboard tube and illuminated from the top with a neon light and one animal was introduced in each jar. After exactly 30 minutes, the animal was removed and the sediment surface was again photographed (time 1), in the same position and under the same light conditions. Using the software ImageJ (http://rsb.info.nih.gov/ij/), the pictures were converted to grey scale and a central circular area was cropped to exclude shadows from the edge of the jar (Fig. 1B). Pictures were processed such that the browsing traces of the animals on the sediment surface resulted black areas against a white background. Then the number of black pixels was quantified. Pictures taken at time 0 were used to correct the values calculated for the browsing traces when irregularities on the sediment surface were detected (i.e. in cases the picture of time 0 was not entirely white). The pixel values were then log-transformed ([log_10_(X+1000)]; 1000 corresponds approximately to the number of pixels of one individual browsing trace). During the assay, four animals were accidentally damaged while handling and were excluded from the analyses. The body lengths of the animals used for behavior analysis were measured after the behavioral assay.

The adjusted intra-class correlation coefficient for the browsing behavior (equivalent to broad sense heritability) was calculated with a linear mixed effect (LMM) model, with treatment as a fixed effect and clone as a random effect (R software package rptR developmental version; (23)). Confidence intervals and statistical significance were calculated using parametric bootstrapping with 5000 iterations and a randomization procedure with 5000 permutations.

### DNA extraction, library preparation and sequencing

The animals assigned to the microbiota analysis were transferred individually from the sediment treatment jars to 40 mL of autoclaved ADaM for about 2 hours to dilute carryover of unattached bacteria. Then, the animals were transferred into 1.5 ml Eppendorf tubes, the ADaM was removed and the tubes were stored at −20 °C until DNA extraction.

DNA was extracted from single animals using a cetyltrimethylammonium bromide (CTAB)-based protocol. The animals were ground with a sterile pestle in 1.5 ml Eppendorf tubes in a 10 mg/ml lysozyme solution and mixed at 850 rpm and 37 °C for 45 minutes. Then, a 20 mg/ml solution of proteinase K was added and the tubes mixed at 850 rpm and 55 °C for 1 hour. After an RNase treatment (20 mg/ml) at room temperature for 10 minutes, a preheated 2X solution of CTAB was added and the tubes mixed at 300 rpm and 65 °C for 1 hour. After two rounds of chloroform isoamyl alcohol (CIA) purification (1 volume CIA; 8 minutes centrifugation at 12,0000 rpm and 15 °C), a solution of sodium acetate 3M pH 5.2 and isopropanol were added to the DNA solution and the tubes were stored overnight at −20 °C. The following day, DNA was purified by two rounds of 70 % ethanol precipitation and suspended in water. The extractions were then incubated at 4 °C overnight and then stored at −20 °C.

All DNA extractions were conducted over a period of 6 days with samples from the different clone by treatment combinations randomly distributed between the days and one reagent-only negative control extraction included every day. DNA from the sediment samples and from one negative control was extracted on a different day using a commercial kit (PowerSoil^®^ DNA Extraction kit; MO BIO Laboratories, cat. 12888-100).

We sequenced amplicons of the V3-V4 variable region of the bacterial 16S rRNA gene using the Illumina MiSeq platform. Amplicons were generated using NEBNext High Fidelity PCR Master Mix (New England Biolabs catalog#M0541L) for 27 cycles in 25 μ1 reactions containing 3% DMSO. The primers used were 341F (5’-TCGTCGGCAGCGTCAGATGTGTATAAGAGACAGGA-3’) and 785R (5’-GTCTCGTGGGCTCGGAGATGTGTATAAGAGACAGCAGA-3’) with Illumina adapter sequences and 0-3 bp random frameshifts. PCR product was purified with Ampure beads at a 0.6x ratio of beads to PCR product, amplified for 9 cycles with Nextera XT v2 indexing primers, and purified again. Libraries were quantified with Qubit and quantitative PCR, normalized, and pooled, followed by additional bead purification to remove remaining short fragments before sequencing on the Illumina MiSeq (reagent kit v3, 300 bp paired-end reads). The same library pool was used for two MiSeq runs; after checking that there was no statistical difference in community composition between the runs (Adonis analysis of Bray-Curtis dissimilarity between samples, p=0.394), the data from the two runs were merged.

### Sequence quality control

Raw reads were quality controlled with FastQC (Babraham Institute, UK). Paired reads were merged (FLASH vl.2.9), primers trimmed (Cutadapt vl.5), and quality filtered (PRINSEQ-lite v0.20.4). OTU clustering including abundance sorting and chimera removal was performed using the UPARSE workflow (24). Only those OTUs represented by 5 or more reads in the run were included. Taxonomic assignment was performed using UTAX against the GreenGenes vl3/5 database. We analyzed samples with more than 5000 total reads. This left 214 samples; numbers of replicates for each combination of variables are reported in Table 1.

Since individual *Daphnia* contain low bacterial biomass, we considered the issue of reagent contamination with bacterial DNA (25). Samples were processed in haphazard order, so erroneous sequences originating from reagent contamination were expected to be distributed randomly and not confounded with any treatment or genotype. For our research question, we were interested in patterns of diversity and changes in composition in response to experimental factors rather than in the presence or absence of any particular strain. For all analyses, we first tested for processing batch effects and stratified the main analysis by batch if they were significant.

For statistical analyses in which host clone was a fixed effect, we excluded clone NO-V-7, since it did not have at least 3 replicates in each treatment; we included this clone in analyses where clone was treated as a random effect. We examined the effects of experimental factors on both overall diversity and the community composition of each animal’s microbiota using standard ecological diversity indices and ordination methods. To evaluate the effect of animal behavior on microbiota, we used as proxies for individual behavior either the mean browsing intensity index of the clone or the browsing intensity of the individual co-housed with the sequenced individual in the same jar (“jar-mate”).

Analyses were carried out in R (3.4.3), using the packages phyloseq (1.22.3), vegan (2.4.6), plyr (1.8.4), dplyr (0.7.4), DESeq2 (1.18.1), nlme (3.1.131), lme4 (1.1.15), metacoder (0.2.0.9012), and ggplot2 (2.2.1).

## Results

### Browsing intensity, animal micrnhinta. and sediment bacteria

Consistently with previous studies (Arbore et al. 2016), browsing behavior intensity varied among *Daphnia* clones (Fig. 2). Clone and treatment, but not their interaction, had a significant effect on browsing behavior (analysis of variance: clone F=12.717, df=11, p<0.0001; treatment F=4.100, df=2, p=0.018; clone*treatment F=1.274, df=22, p=0.193). The average browsing intensity of animals from the NET treatment was lower than that of animals in the SED and AUT treatments (Fig. S1). The total phenotypic variance for browsing behavior explained by clone, after controlling for the treatment effect, corresponded to 36.5% (95% Cl = [13.2, 55.3%], p = 0.0002). Clone but not treatment had a significant effect on body size, so we assume that access to (and type of) sediment did not substantially affect nutrition and growth over the timeframe of the experiment (analysis of variance: clone F=8.08, df=11, p < 0.001; treatment F=2.01, df=2, p=0.137; clone*treatment F=1.43, df=22, p=0.103). Individual body size was uncorrelated with behavior (analysis of variance: F=0.346, df=1, p=0.56; Fig. S2).

**Figure 2:**
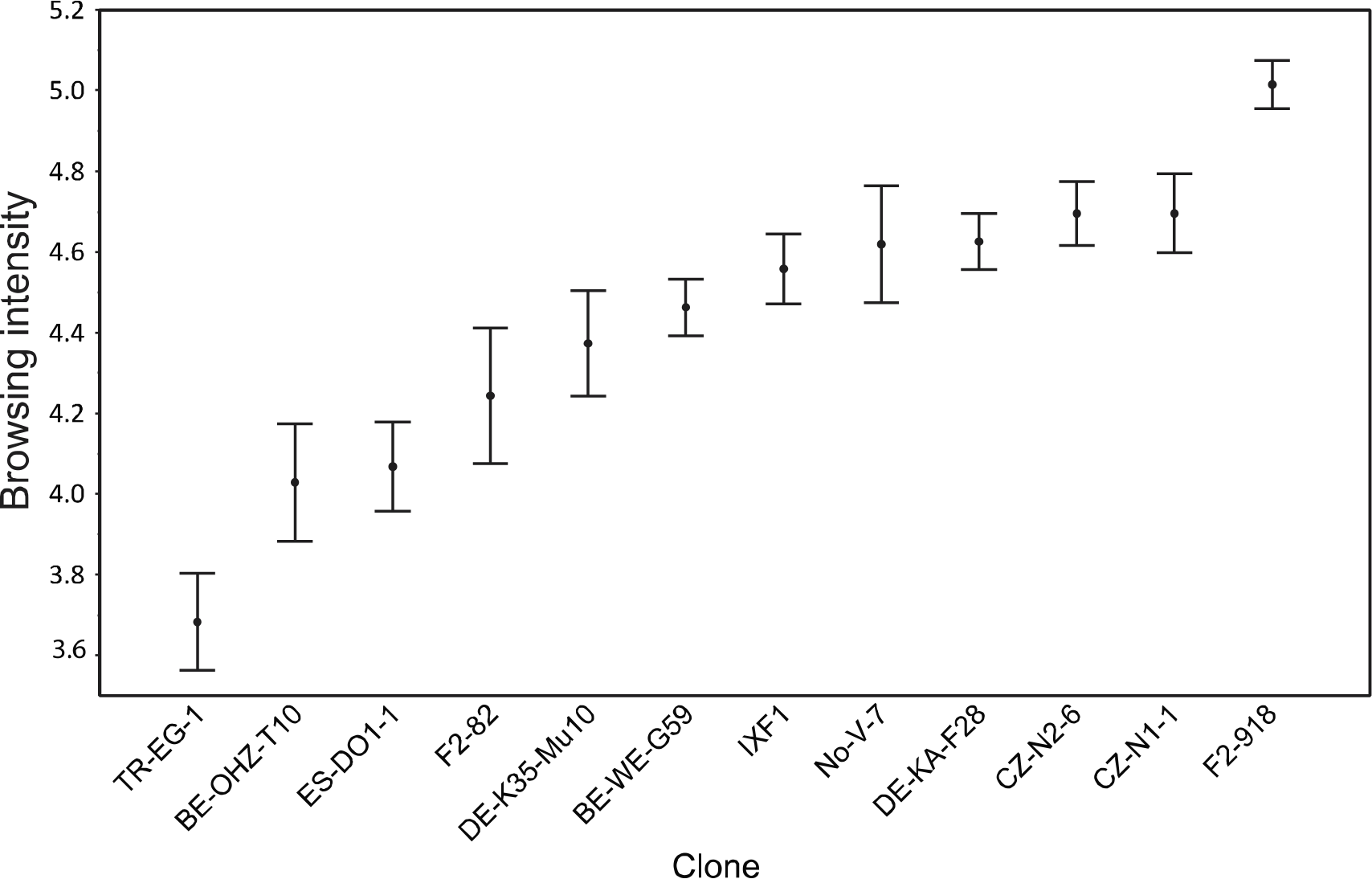
Browsing intensity of 12 *D. magna* clones (mean and SE). Browsing intensity was defined as the Logio of the area of the browsing traces left by individual replicate animals browsing on a sediment surface for 30 minutes (see Fig. 1).

A total of 370 OTUs were found among the animal samples; of these, 318 were found in less than 10% of samples. (See Fig. S3A-C for taxonomic heat trees of OTUs with presence/absence information) (26). Consistently with multiple previous studies of *Daphnia* microbiota (27–30), the most abundant bacterial species was a single strain (OTU_1) of *Limnohabitans* sp (Betaproteobacteria, Comamonadaceae), with a mean relative abundance across all clones of 0.39 (s.e.m. 0.02). Interestingly, a second strain of *Limnohabitans* (OTU_2) was a dominant strain only in the three clones originating from clones bred in the laboratory as part of a genetic breeding design (QTL panel; 0.32 mean relative abundance among individuals of clones IXF1, F2-82, F2-918; 0.0016 mean relative abundance in remaining clones). As expected, the sediment originating from the SED treatment had much higher bacterial species richness than that originating from the AUT treatment (Fig. S4).

### Effects of treatment and clone on alpha diversity

Both *Daphnia* clone and treatment, but not their interaction, had significant effect on the Shannon and inverse Simpson alpha diversity indices (Table 2). For further analyses, we focused on the Shannon index, because it takes into account not only species richness but also evenness (with additional species given more weight as they become more abundant). The Shannon index displayed no significant effect of processing batch (df=5, F=1.42, p=0.22). Shannon diversity estimates for the 12 clones arranged in order of increasing average browsing intensity and the three groups (AUT, NET and SED) are shown in Fig. 3 (species richness and inverse Simpson index are shown in Fig. S5A-B). Unexpectedly, the highest average alpha diversity in most clones (9/12) was observed in the AUT treatment group, despite their exposure to less-diverse sediment than the SED group. Therefore, diversity of animal microbiota does not directly reflect diversity of bacteria in the environment.

**Table 2.** Results of analyses of variance of different alpha diversity indices. All treatments are included; clone NO-V-7 is excluded.

|  | Richness | Shannon | Inverse Simpson |
| --- | --- | --- | --- |
| Clone | F=0.40, df=10,<br>**p=.944** | F=2.43, df=10,<br>**P=.00997 \*** | F=1.98, df=10,<br>**p=.0383 \*** |
| Treatment | F=4.91, df=2,<br>**p=.00842 \*** | F=12.21, df=2,<br>**p<.0001 \*** | F=13.25, df=2,<br>**p<.0001 \*** |
| Clone:Treatment | F=0.75, df=20,<br>**p=.770** | F=0.78, df=20,<br>**P=.734** | F=0.65, df=20,<br>**P=.8124** |

**Figure 3:**
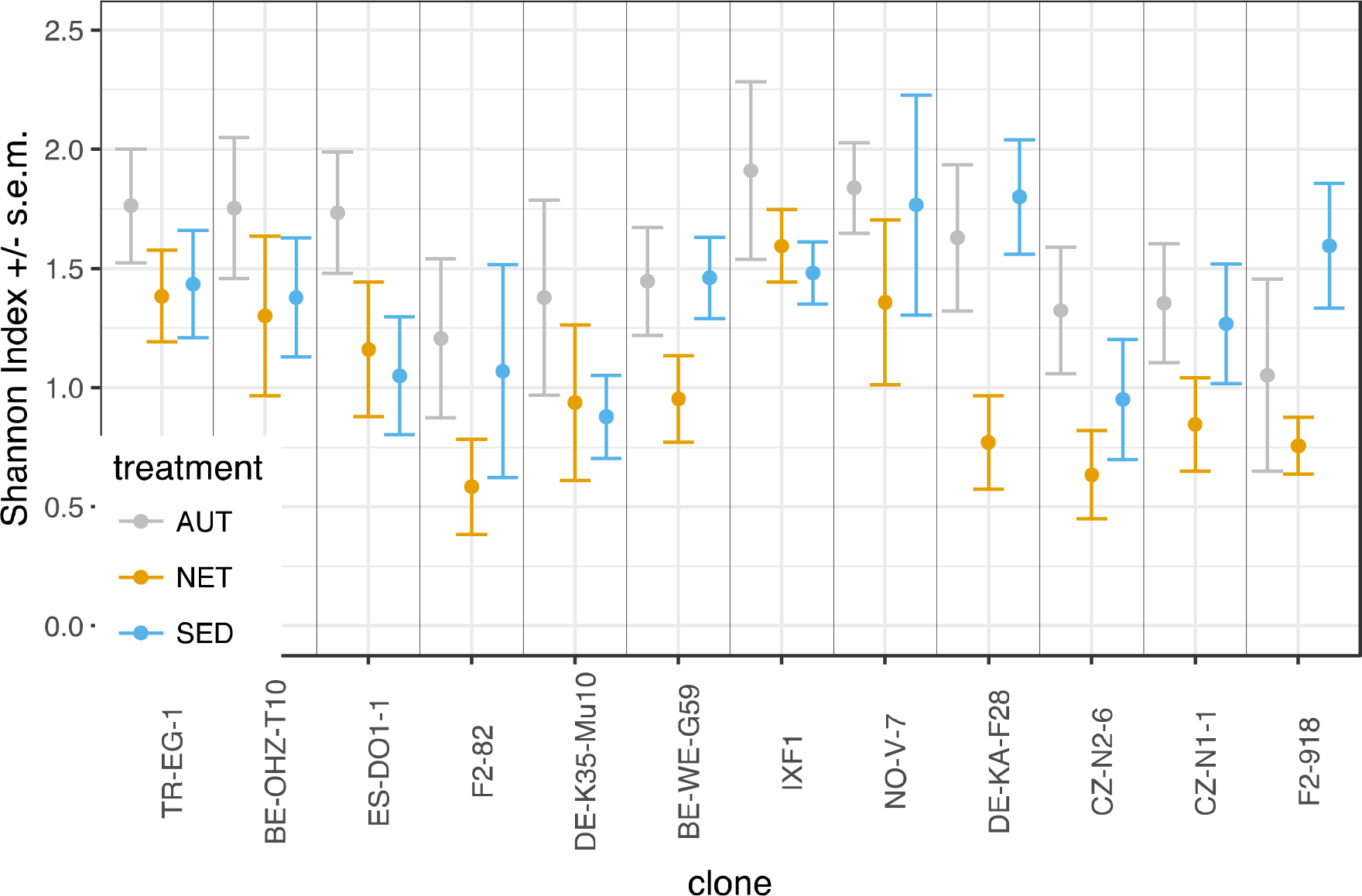
Microbiota diversity (Shannon index) of *Daphnia* clones under three different treatment conditions (AUT, NET and SED). AUT: Exposure to autoclaved sediment; NET: prevented exposure to untreated sediment; SED: exposure to untreated sediments. Error bars represent standard error of the mean. Clones are arranged left-right by increasing average clone browsing intensity.

To specifically investigate the effect of direct access to the same bacteria-rich sediment, we compared the NET and SED treatment groups’ diversity as a function of clonal average behavior in each group (Fig. 4A). The difference in mean Shannon diversity between SED and NET animals was greatest at the highest average clonal level of browsing intensity (Fig. 4B; linear regression p=0.055). A similar tendency could be seen when the browsing intensity of each individual’s jar-mate was used as the proxy for individual behavior (Fig. S6). Shannon diversity significantly depended on the interaction between treatment and clonal average browsing intensity in a linear mixed-effects model with clone included as a random effect (Table 3); the same was true when treatment-specific clonal average behavior was used as the behavior proxy, but not when individual jar-mate behavior was used (Table S1).

**Figure 4:**
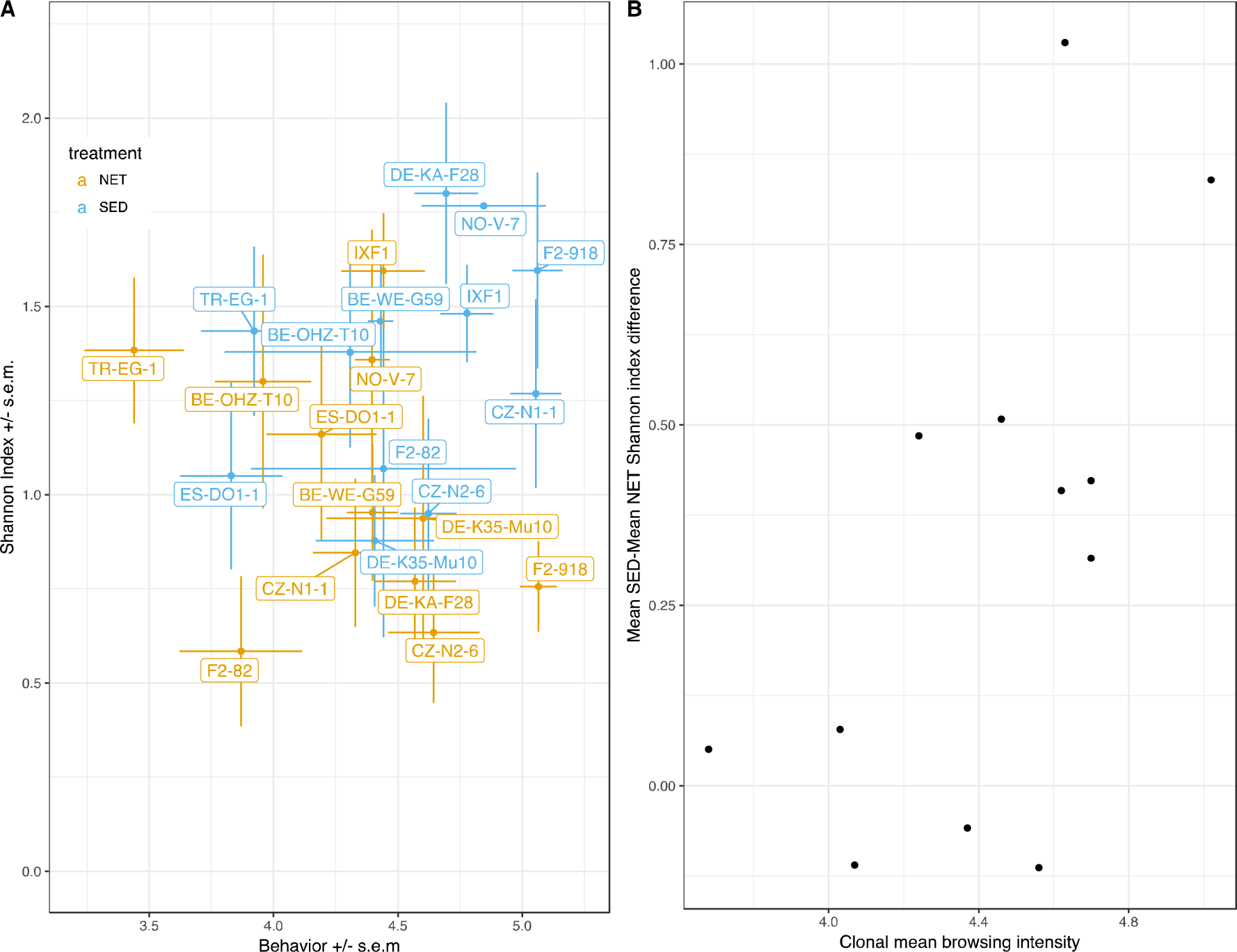
Average browsing intensity and average microbiota diversity in the NET and SED treatments. A: average clone browsing intensity and average clone microbiota diversity in the NET and SED treatments. Average browsing intensities were calculated based on samples whose jar-mates passed the sequence quality control (N=214, Table 1). B: average clone browsing intensity and the difference between average Shannon diversity in the SED treatment and average diversity in the NET treatment. Here, average browsing intensities were calculated based on the complete set of samples (N=228, i.e. all assayed jar-mates). Error bars represent standard error of the mean.

**Table 3.** Effect on Shannon index. NET and SED treatments only, all clones included. Linear mixed-effects model with treatment, clonal average browsing intensity and clonal average size as fixed effects and clone as random effect.

|  | numDF | denDf | F-value | p-value |
| --- | --- | --- | --- | --- |
| (Intercept) | 1 | 129 | 270.12 | <.0001 |
| **Treatment** | **1** | **129** | **12.26** | **0.0006 \*** |
| Clone average behavior | 1 | 9 | 0.275 | 0.613 |
| Clone average size | 1 | 9 | 2.568 | 0.144 |
| **Treatment:Clone average behavior** | **1** | **129** | **4.677** | **0.0324 \*** |
| Treatment:Clone average size | 1 | 129 | 0.0980 | 0.755 |

### Community composition and acquisition of bacteria from sediment

To examine shifts in bacterial community composition in response to environmental treatments, we Hellinger-transformed the bacterial abundances by taking the square root of the relative abundance of each taxon in each sample to reduce the influence of rare taxa, and then calculated pairwise Bray-Curtis distances between samples. The average distance to the centroid (dispersion) was lower in the NET group than in the AUT and the SED groups (Fig. 5), suggesting that access to sediment increases variability of microbiota regardless of the composition of the sediment. To see whether the different sediments resulted in systematically different microbiota composition, we excluded the NET group and performed principal coordinates analysis (PCoA) (Fig.6). Adonis analysis stratified by processing batch showed that both treatment and clone had a significant effect (treatment: R^2^=0.05, p=0.001; clone: R^2^=0.16, p=0.001), but not their interactions (treatmentxlone p=0.63). However, clones also showed significant differences in dispersion (p<0.001).

**Figure 5:**
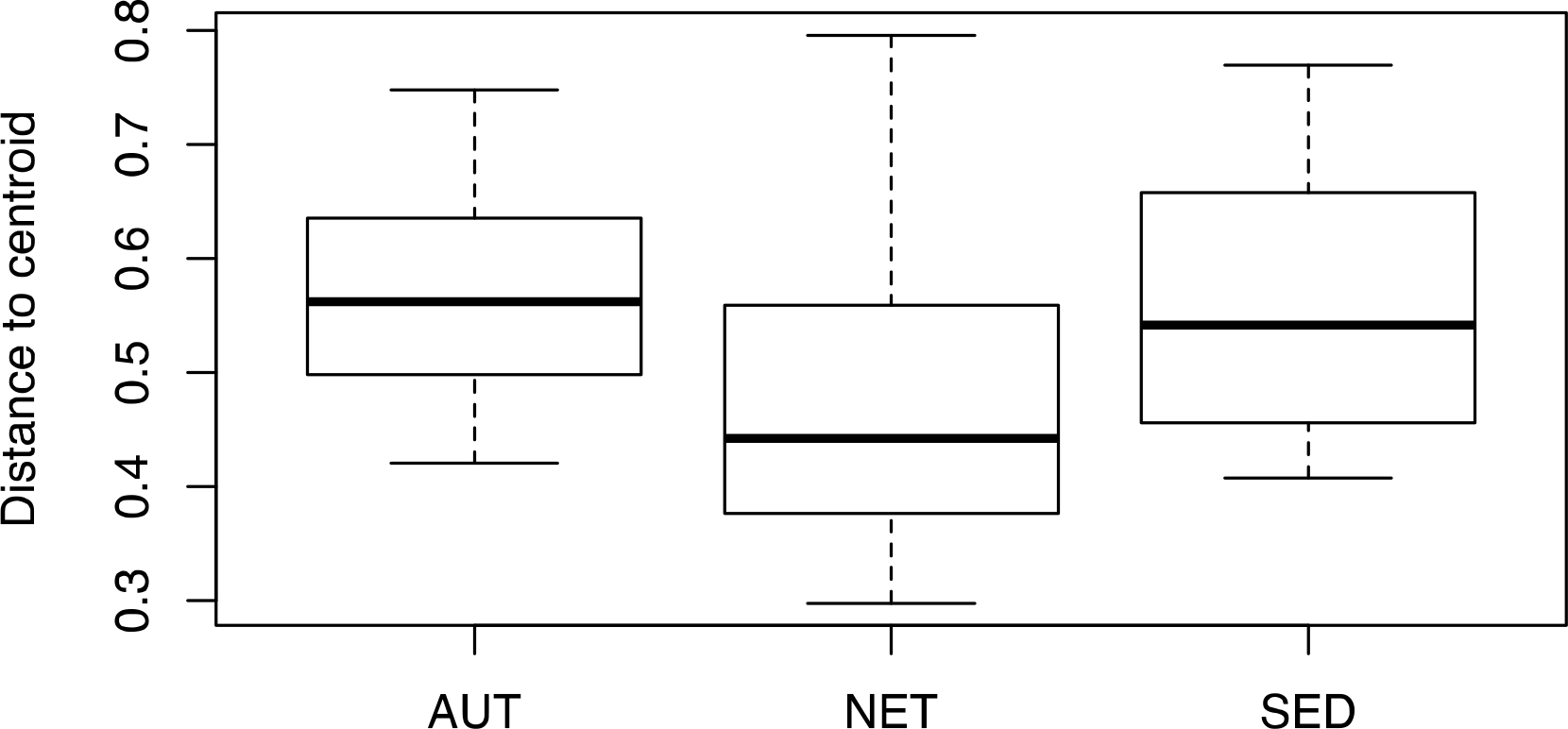
Within-group dispersion of community similarity. The median distance to the centroid is lower in the NET treatment group than in the others (Permutation test of multivariate dispersion p<0.0001, 999 permutations), meaning that NET communities are less variable than AUT or SED microbiotas.

**Figure 6:**
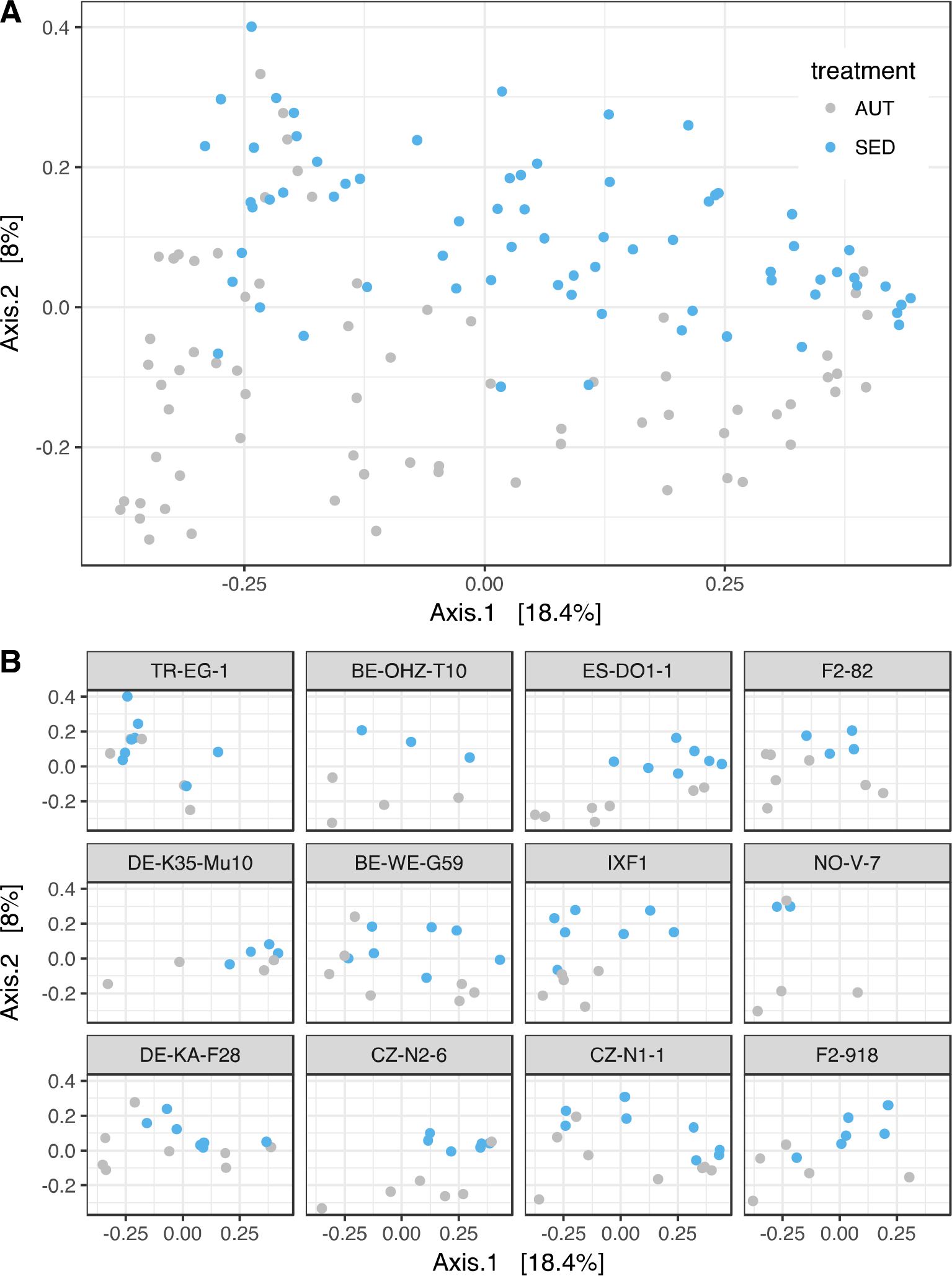
Similarity of bacterial community composition in the AUT and SED treatments. A: first and second axis of a principal coordinates analysis (PCoA) of bacterial community composition based on Hellinger-transformed Bray-Curtis dissimilarities. B: first and second axis of a principal coordinates analysis (PCoA) of bacterial community composition by *Daphnia* clone.

Having confirmed that differences in the sediment environment resulted in differences in animal microbiota composition, we next explored the extent to which environment-specific bacteria contributed to these differences. We used DESeq2 to determine which bacteria were significantly more present in natural sediment than autoclaved sediment (n=3 each). 115 OTUs were calculated to be significantly differentially present between the two sediment types; of these, 48 had at a log2-fold increase of at least 8 in natural sediment compared to autoclaved sediment. We refer to these as natural-sediment-derived taxa. The 8-fold threshold was chosen based on inspecting the data; similar results were seen when sediment-derived bacteria were defined by a log2-fold change of 5 or 10; see Fig. S7. Only one of these OTUs was found in a majority of animals, and the median number of animals in which a given OTU was found was 6.5. We therefore concluded that animals likely acquired environmental bacteria randomly rather than selectively from the environment. Accordingly, we looked at the total relative abundance of reads from all natural sediment-derived bacteria in each individual.

The relative abundance of natural-sediment-derived bacteria was generally low in both the AUT and NET treatment groups, and increased with browsing intensity in the SED treatment group (Fig. 7), with a significant interaction effect between treatment and clonal average browsing intensity (Table 4). Treatment-specific clonal average behavior showed the same significant interaction effect with treatment, but the interaction effect was not significant when jar-mate behavior was used as the behavior proxy. Among the set of clones examined here, an appreciably high relative abundance of sediment-derived bacteria was detectable mainly in clones with a browsing intensity index higher than mean 4.4 (clones IXF1, NO-V-7, DE-KA-F28, CZ-N2-6, CZ-N1-1, F2-918). The mean relative abundance of sediment-derived bacteria in the SED treatment in the pooled animals from these clones was 0. 14 (s.e.m. 0.026), whereas it was 0.048 (s.e.m. 0.0097) in the lower-browsing clones. Across all clones in the AUT and NET treatment groups, the average relative abundance of sediment-derived bacteria was 0.051, nearly identical to that of the low-browsing clones in the SED conditions.

**Figure 7:**
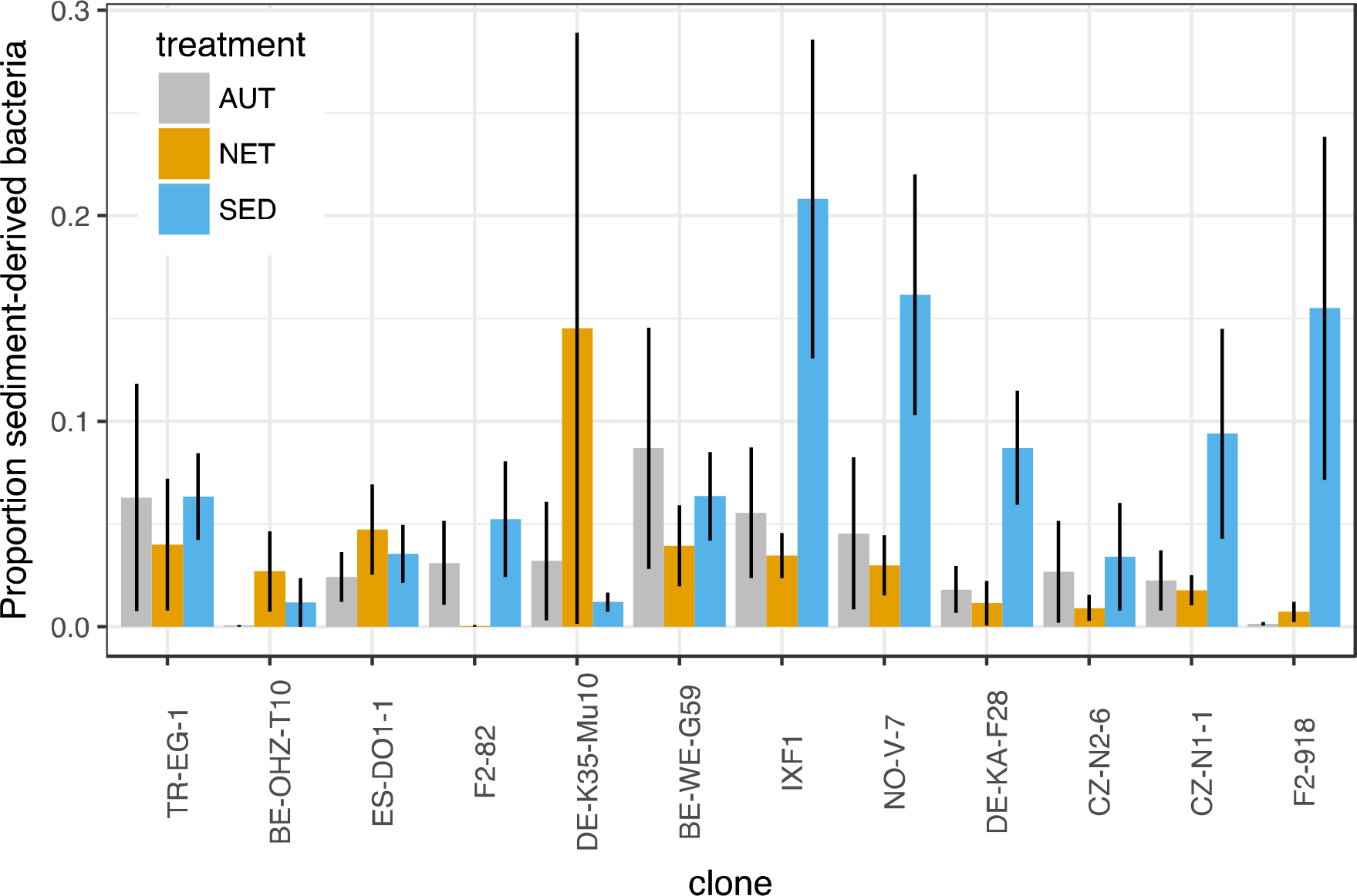
Analysis of sediment-derived bacteria. Proportion of sediment-derived bacteria in the microbiota of animals from AUT, NET and SED treatments. Sediment-derived bacteria were identified by comparing autoclaved and untreated sediment samples (log2-fold increase of at least 8 in natural sediment compared to autoclaved sediment). Error bars correspond to standard errors of the mean. Clones are arranged left-right by increasing average clone browsing intensity.

**Table 4.** Effect on relative abundance of sediment-derived bacteria. All treatments and all animals included. Linear mixed-effects model with treatment and clonal average browsing intensity as fixed effects and clone as random effect.

|  | numDF | denDf | F-value | p-value |
| --- | --- | --- | --- | --- |
| (Intercept) | 1 | 198 | 53.999 | <.0001 |
| **Treatment** | **2** | **198** | **6.500** | **0.0018\*** |
| Clone average behavior | 1 | 10 | 0.490 | 0.4998 |
| **Treatment:Clone average behavior** | **2** | **198** | **4.185** | **0.0166\*** |

## Discussion

Our results have several implications for studies of animal-associated microbiota in diverse environmental settings. First, we confirm the intuition that environmental sources of bacteria affect the diversity of animal microbiota, but not because more diverse environments always create more diverse microbiota; rather, the animals we exposed to the less species-rich autoclaved sediments had higher overall diversity in their microbiota than those exposed to untreated, bacterial-species-rich sediment. We hypothesize that this might be due to competitive interactions between *Daphnia* microbiota and the particular microbes found in these sediments. The untreated sediments may contain bacteria that can outcompete multiple strains of “native,” preexisting *Daphnia* microbiota. If this were the case, then browsing in sediment could have multiple opposing effects on overall microbiota diversity: on the one hand, it would bring daphnids into contact with more diverse bacteria, but on the other hand those bacteria could reduce existing microbiota diversity. In the NET treatment, animals might be exposed to some sediment-derived bacteria in the water column but lack access to the full diversity of bacteria in the sediment. An experiment designed to explicitly test this hypothesis would be required to determine whether there are competitive interactions between exogenous sediment-derived bacteria and those typically carried by *Daphnia* in the laboratory; it would also be interesting to see how these competitive effects interact with early colonization events in young *Daphnia*.

We also saw that having access to either sediment increased the variability of community composition as measured by multivariate dispersion. These results suggest that having access to multiple habitats with different bacterial communities can affect the diversity and composition of an animal’s microbiota. Therefore, fine-scale heterogeneity in a host’s habitat might be a relevant aspect to take into account when examining effects of environment on animal microbiota. This is especially important when considering ecological immunology, because disease-causing bacteria in the environment may cause short-term risk but also long-term fitness benefit via processes like immune priming (31, 32).

Our data further suggest that the diversity of *Daphnia*-associated microbiota in a particular environment may to some extent be mediated by genotype-specific sediment browsing intensity. This was apparent as the net barrier made the greatest difference in microbial alpha diversity in high-browsing host clones. However, this effect may be partially obscured by several factors: the hypothesized competitive exclusion effects we allude to above, and also non-behavior-related host genotype effects on microbiota diversity. While host genotype had an effect on microbial diversity, the highest- and lowest-browsing clones in our study had similar microbial alpha diversity overall. The only way to conclusively determine that differences between the microbiotas of different genotypes are mediated by host behavior independently of other host traits would be to genetically manipulate behavior on an otherwise identical genetic background; we approximate this in our experiment with the treatment where *Daphnia* are blocked from sediment browsing, contrasted with the treatment where they are allowed to browse freely. Our cautious conclusions about the effect of behavior on microbiome are based on examining the contrast between these treatments within each genotype, not based on the observation of genotype-dependent differences alone. It was only in evaluating the difference between presence and absence of the barrier that an effect of browsing on diversity could be seen. We conclude that the effect of environmental bacteria on host-associated microbiota is not additive. The clearest effect of environmental bacteria on host-associated microbiota was not on alpha diversity, but relative abundances of certain taxa

Clones with low average browsing intensity had no greater amount of sediment-specific bacteria than animals exposed to autoclaved sediment or prevented from browsing, whereas those with high browsing intensity could reach over 60% of reads from environment-derived bacteria in some individuals. While many studies of animal microbiota rightly concern themselves with distinguishing between truly “host-associated” microbiota versus “transient environmental” microbiota, these results raise the possibility that the amount of environmental microbes found in association with an animal could itself be a host-genotype-specific feature of the microbiome.

Another key question is whether browsing behavior affects community composition by simple exposure to more colonizing bacteria, or by more frequent replenishment of bacterial taxa that would not otherwise persist in association with the host (33). For example, browsing frequently enough may replenish bacteria that would otherwise be lost when the animal molts. In this study, we made no assumptions about the type of the interactions between the sediment-associated bacteria and the *Daphnia*, but still were able to see that presence of sediment-associated bacteria affected the bacteria that *Daphnia* were carrying.

It would also be interesting to investigate whether carriage of bacteria on *Daphnia* from the sediment into the water column affects bacterial dynamics in the larger environment; previous studies have shown that movement of *Daphnia* between benthic and limnetic environments represents a mechanism of bacterial dispersal in the environment (34). Studies using classification methods more sensitive than 16S-based taxonomy may be necessary to unambiguously distinguish and assign sources to different bacterial strains.

## Conclusion

We show that at least some characteristics of host-associated microbial community composition result from genotype-by-microhabitat interactions, specifically ones resulting from genotype-specific variation in behavior. We show this using an experimental treatment that externally manipulated behavior, but genetically manipulating behavior to confirm these results would be a natural next step when the molecular tools to do so become available. Behavior could thus be considered a genetic factor that shapes microbial exposure in a given environment. Overall, these results provide further evidence that environment, behavior, genetics, disease risk, and microbial community composition are interrelated in potentially complex ways. Our observations indicate a need for more integrative eco-immunology studies, in which the interfaces between behavioral ecology, microbial community ecology and evolution of immune function are explored. Studies can take advantage of the experimental tractability of the *Daphnia*-microbiota system to further investigate these relationships in mechanistic detail.

## Acknowledgments

We thank Daniel Lüscher for designing the experimental jars, Jürgen Hottinger for laboratory support, and the Botanical Institute of the University of Basel for providing sediment. Illumina sequencing was performed at the Genetic Diversity Centre of the ETH Zürich; thank you to Aria Minder and Silvia Kobel for advice on library preparation.

## Author contributions

AAM and RA conceived the study and performed the experiment with input from DE. RA performed behavioral assays and analysis. AAM and RA prepared sequencing libraries. JCW performed quality control and OTU clustering of sequence data. AAM analyzed sequencing data. AAM and RA wrote the paper. DE and JCW revised the paper.

## Data deposition

Sequence data will be deposited in the European Nucleotide Archive of the EBI
upon publication. Data tables and code used for analysis can be found on Github at
https://github.com/amusheg/Daphnia-microbiota-behavior and will be deposited in Dryad upon publication.

